# HuConTest: Testing human contamination in great ape samples

**DOI:** 10.1101/2021.03.30.437753

**Authors:** Martin Kuhlwilm, Claudia Fontsere, Sojung Han, Marina Alvarez-Estape, Tomas Marques-Bonet

**Affiliations:** Institut de Biologia Evolutiva, (CSIC-Universitat Pompeu Fabra), PRBB, Doctor Aiguader 88, Barcelona, Catalonia 08003, Spain; CNAG-CRG, Centre for Genomic Regulation (CRG), Barcelona Institute of Science and Technology (BIST), Baldiri i Reixac 4, 08028 Barcelona, Spain; Institucio Catalana de Recerca i Estudis Avançats (ICREA), Barcelona, Catalonia 08010, Spain; Institut Català de Paleontologia Miquel Crusafont, Universitat Autònoma de Barcelona, Edifici ICTA-ICP, c/ Columnes s/n, 08193 Cerdanyola del Vallès, Barcelona, Spain

**Keywords:** Contamination, non-human primates, next generation sequencing, fecal DNA, ancient DNA

## Abstract

Modern human contamination is a common problem in ancient DNA studies. We provide evidence that this issue is also present in studies in great apes, which are our closest living relatives, for example in non-invasive samples. Here, we present a simple method to detect human contamination in short read sequencing data from different species. We demonstrate its feasibility using blood and tissue samples from these species. This test is particularly useful for more complex samples (such as museum and non-invasive samples) which have smaller amounts of endogenous DNA, as we show here.

**Significance statement:** Human contamination can be a confounding factor in genomic studies, especially in the case of fecal, museum or ancient DNA from great apes. It is important for quality assessment, screening purposes and prioritization to identify and quantify such contamination. The tool presented here is a simple and versatile method for this purpose, and can be applied to a wide range of sample types.

## Main Text

Contamination from exogenous sources is a problem common in ancient DNA, where multiple tools exist (Peyrégne & Prüfer 2020), as well as in studies of non-human primates (Prado-Martinez et al. 2013). Specifically, human contamination may occur in great ape samples of various origin and quality. Previously, differences in the mitochondrial genome between species were used to assess contamination (Prado-Martinez et al. 2013), which is a sensible strategy for high-coverage data. However, this approach is of limited use for shallow shotgun sequencing, especially of samples with low endogenous DNA content, such as fecal, historical, or ancient samples, as well as sequencing data obtained after enrichment through capture (Fontsere et al. 2020). Here, we devise a strategy based on diagnostic sites dispersed across the autosomes which can help detecting human contamination in an unbiased manner and with sparse data available.

### Determination of diagnostic sites

We used previously published diversity data on high-coverage genomes from all great apes and modern humans (Table S1, Figure 1A), specifically, genomes from 58 chimpanzees and 10 bonobos (*Pan* clade) (Prado-Martinez et al. 2013; De Manuel et al. 2016), 43 gorillas (*Gorilla* clade) (Prado-Martinez et al. 2013; Xue et al. 2015), 27 orangutans (*Pongo* clade) (Prado-Martinez et al. 2013; Nater et al. 2017) and 19 modern humans (Mallick et al. 2016). All genomes were processed as described previously (De Manuel et al. 2016): Sequencing data was mapped to the human genome (hg19) using BWA-MEM 0.7.7 (Li & Durbin 2009), PCR duplicates were removed using samtools (Li et al. 2009), and reads were locally realigned around indels using the GATK IndelRealigner 3.4-46 (McKenna et al. 2010). Genotypes were obtained individually using GATK UnifiedGenotyper with the EMIT_ALL_SITES parameter, and GVCFs from individuals were merged with GATK CombineVariants. The three species complexes *Pan*, *Gorilla* and *Pongo* were then filtered separately: Biallelic SNPs within each species complex together with humans were retrieved, and filtered to exclude repetitive regions of the genome and regions with low mappability (35mer mappability). Finally, for each individual, genotypes were set to missing at sequencing coverage lower than 6 and higher than 100, and with a mapping quality lower than 20.

**Fig. 1.**
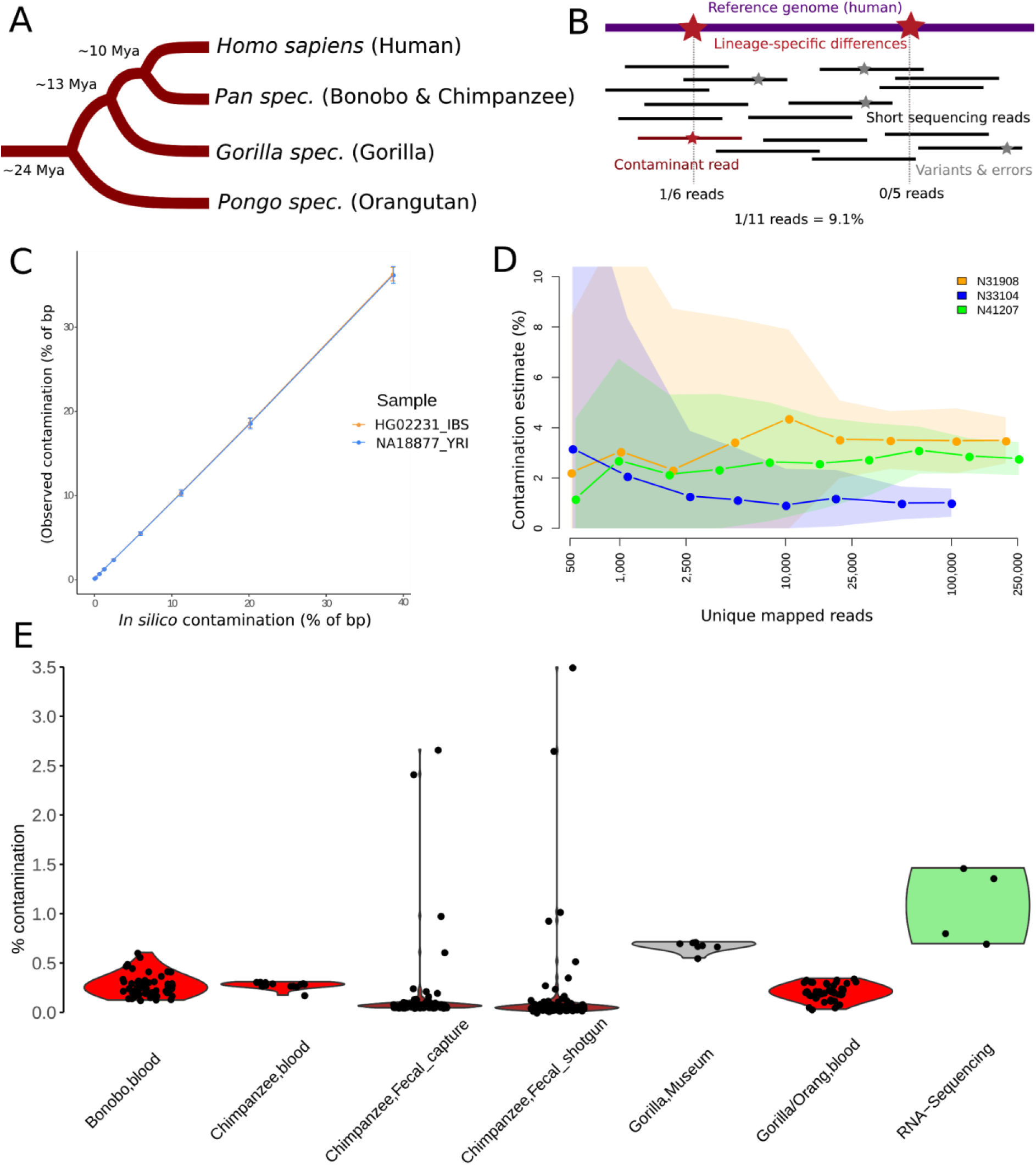
Summary of this study. A) Schematic tree of the great ape species, with approximate divergence times (Besenbacher et al. 2019). B) Schematic representation of the method. C) Performance of detection of *in silico*-contamination in a gradient from ~0.1-39%, point estimate ± one standard deviation. D) Performance when downsampling sequencing data from fecal samples with 1-3.5% of human contamination. Point estimates ± one standard deviation. E) Contamination estimates of blood samples for sequencing libraries from all species (red; bonobo N=52, chimpanzee N=15; gorilla/orangutan N=47), fecal samples before and after capture (brown; N=109, without sample N42003 which has high levels of non-great ape contamination), museum samples (grey; N=8) and RNA sequencing data (green; N=4).

We retrieved SNPs where at least 98% of the chromosomes in the species complex showed the alternative allele (different from the human reference), with less than 5% of missing genotypes, and where all modern human chromosomes included in this study carried the human reference allele, without allowing for missing genotypes. We allowed for residual amounts of human-reference-like genotypes in the great ape species, to account for residual sources of error in the reference set that might result in erroneous rare variation, and we deemed tolerating these a conservative strategy for determining diagnostic sites. Across the whole genome, we find 4,460,987 diagnostic sites for *Pan* species, 6,981,108 diagnostic sites for *Gorilla* species, and 7,518,570 diagnostic sites for *Pongo* species. The differences between species are partially explained by their evolutionary divergence to humans (Kaessmann & Pääbo 2002; Kuhlwilm et al. 2016; Prado-Martinez et al. 2013), but also the number of individuals used, as well as sequencing quality and coverage. We used the R package rtracklayer (Lawrence et al. 2009) to perform a liftover of these positions to the human genome version 38 (GRCh38).

### Contamination assessment and performance

Contamination is defined here as the proportion of observed human reference alleles at diagnostic positions in short sequencing reads (Figure 1B). The test itself is wrapped in an R script (confirmed to work with R versions 3.2.0, 3.4.4, 3.5.0, 3.6.0, and 4.0.1 (R Core Team 2015)), to directly process the number of reads carrying the reference or the alternative allele. We use samtools mpileup (tested for version 1.0 and 1.9) to obtain read depth and alternative read depth at diagnostic sites, and join these data with information on the alternative allele in the test species. We then calculate the number of reads matching the target species complex allele, and subtract this value from the total read depth, thus retrieving the number of reads matching the human reference allele (more strictly, not matching the target species allele). We perform this calculation for each chromosome separately in order to obtain the standard variation, and report the genome-wide point estimate, one standard deviation, and the number of positions observed by the test. The latter value is useful to assess the reliability of the test at extremely shallow sequencing. The test can be applied to files with a bam or cram extension, containing short sequencing reads mapped to the human genome (hg19 or GRCh38). The basic filtering at this step can be simple, but it is advisable to remove adapter sequences (Schubert et al. 2016) and PCR duplicates to assess the unique contaminant fraction, as well as unmapped reads, non-primary alignments and sequences with a low mapping quality (<30). We specifically recommend filtering the sequences on fragment/insert length to avoid spurious alignments, which may happen at a high rate in the case of samples with large amounts of bacterial DNA (Meyer et al. 2016).

We tested the contamination test by artificially introducing modern human sequencing reads into bam files from the other species (*in silico* contamination), using eight human individuals that were not part of the reference panel (Table S2) (Auton et al. 2015), and great ape samples from other studies (Locke et al. 2011; Prüfer et al. 2012; Besenbacher et al. 2019). First, each human bam file was downsampled to ~1.14M reads and merged with a chimpanzee bam file (ERR032960), to simulate ~5% of human contamination. Since the read length differs between sequencing libraries from different studies, we account for the expected amount of human contamination by using the percentage of human base pairs added to the final bam file. After running the human contamination test in each file, we detect an average of 5.5% human contamination (Table S2), with minimal differences between humans from different world regions. When testing a gradient of increasing amounts of introduced human sequences from ~0.1% to ~39% to a chimpanzee bam file (Table S3, Figure 1C), the contamination is estimated correctly. The test is performing well for *in silico* contamination from modern humans in each of the great ape species (Table S4).

We also determined the inferred amount in the case of cross-testing, i.e. performing the test of species-specific sites from other species (Table S5). Here, we find estimates of 44-82% attributed to contamination, depending on the species combination, which is a consequence of the shared ancestry between humans and the other species. This demonstrates that the test is species-specific, and large amounts of reads that do not carry species-specific alleles will be detected when a different primate species is present.

### Application to other sample types

We first applied the test to blood samples from all great ape species, which are generally expected to contain at most small amounts of human contamination. For 67 randomly chosen sequencing libraries from seven chimpanzee and four bonobo individuals (Prüfer et al. 2012), we found an average of 0.28% (0.13-0.61%) of reads that are putatively due to human contamination (Figure 1E). Four tissue samples from chimpanzees (White et al. 2019) show low estimates of contamination (0.03-0.067%), as expected for samples likely not containing true contamination. Similar results are obtained for four libraries from gorilla (0.033-0.159%, on average 0.075%) and 43 libraries from orangutan (0.08-0.35%, on average 0.22%) blood samples (Besenbacher et al. 2019; Locke et al. 2011). We conclude that traces of putative human contamination are observed, if at all, only at very small amounts in sequencing data from great ape blood samples. These estimates are conservative, since sequencing errors, mapping reference bias and variation in these individuals may contribute to these numbers, especially considering that error rates of these sequencing technologies were decreasing after the publication of some of these studies (Prüfer et al. 2012; Locke et al. 2011). We also note that results for data mapped to hg19 and hg38 are almost identical (Table S6).

We then applied the contamination test to non-invasive samples which usually contain small amounts of host DNA, and may require target hybridization methods to obtain sufficient data (Hernandez-Rodriguez et al. 2017; Fontsere et al. 2020). We applied our method to shotgun and exome capture sequencing data that were obtained from the same 109 sequencing libraries from chimpanzee fecal samples (White et al. 2019). We found an average of 0.35% (0-24.6%) human contamination for the pre-capture (shotgun) and 0.32% (0.05-21%) human contamination in the post-capture (enriched) sequencing data (Table S6, Figure 1E), with strong correlation for the same samples (r=0.99, p-value < 2.2×10-16). We find one sample with an estimate of 24.6% and three more samples with more than 1% of human contamination (Table S6). In case of fecal samples collected from the field that may contain other mammalian DNA than the target species through diet or mis-identification, it is advisable to perform a competitive mapping of sequences when large amounts of contamination are detected. This will help to determine the species of origin, for example using BBSplit (https://sourceforge.net/projects/bbmap/) with a reference panel of great apes, and possibly other primate species living in the same habitat. We applied this method to these four samples (N42003_Shotgun1, N31908_Shotgun1, N33104_Shotgun1 and N41207_Shotgun1), and find that the main contaminant in one sample is most likely another primate rather than human (Table S7). It is known that chimpanzees hunt other primates (Boesch & Boesch 1989), and DNA from primate prey can persist in the feces. We conclude that the design of the contamination test presented here is able to identify reads carrying mutations that differ from the target species, even if these are not human-specific. When applying BBsplit method to *in silico*-contaminated samples, we confirm humans as the source of the contamination – although with less precision regarding the amount of contamination when compared to our method – while the majority of unambiguously mapped sequences align to the target species (Table S7).

Our analysis shows that DNA extracts/libraries from fecal samples are occasionally contaminated, and may need to be removed from certain downstream analyses. Hence, it is advisable to perform a contamination test for sequencing data from this type of sample, comparable to ancient and historical samples. We assessed the power to detect human contamination with very shallow sequencing, by downsampling the sequencing reads of the three fecal samples from White *et al*. (N31908_Shotgun1, N33104_Shotgun1 and N41207_Shotgun1) with 1-3.5% human contamination. We downsampled these in several steps down to ~1,000 production reads (Table S8), and calculated the estimated amount of human contamination. These results (Figure 1D) confirm that our method is robust in confidently detecting human contamination even in the case of very shallow sequencing, as low as ~1,000 reads aligned to the human reference genome, although with high standard deviation. In the case of fecal samples with around 5% of estimated hDNA, this could be as little as ~20,000 production reads, making the test applicable to shallow data from an initial screening procedure (Fontsere et al. 2020).

We also applied the test to published sequencing data from eight museum samples from gorillas (van der Valk et al. 2019). Here, we find an estimated contamination of on average 0.68% (0.55-0.72%), which is slightly lower than the reported estimates which were based on mitochondrial diagnostic loci (0.28-1.67%, on average 1%), and slightly higher than estimates for blood samples, as expected for museum specimens that have been handled by humans. Contamination estimates from mitochondrial and nuclear loci from the same sample have been found to not be identical in hominin samples (Prüfer et al. 2014), and at shallow sequencing coverage a small number of reads would overlap with mitochondrial diagnostic loci. Still, the differences between these methods are minor, and results on data mapped to hg19 and hg38 are almost identical (Table S6), as is the case for blood samples. Finally, we performed the contamination test on RNA-sequencing data from great ape tissue samples (Brawand et al. 2011), mapped using tophat2 (Kim et al. 2013). We find slightly higher amounts of contamination (Table S6), either due to real contamination in the samples, or to higher error rates and mapping bias in transcriptome data compared to genome sequencing data.

## Method availability

The contamination test script including documentation is publicly available on GitHub: https://github.com/kuhlwilm/HuConTest. Files with the diagnostic positions are publicly available on FigShare (doi: 10.6084/m9.figshare.14237834).

## Data availability

There are no new data associated with this article.

## Acknowledgments

We thank Marc de Manuel for help with data preparation. M.K. is supported by “la Caixa” Foundation (ID 100010434), fellowship code LCF/BQ/PR19/11700002. M.A.E. is supported by an FPI (Formación de Personal Investigador) PRE2018-083966 from Ministerio de Ciencia, Universidades e Investigación. T.M.-B is supported by funding from the European Research Council (ERC) under the European Union’s Horizon 2020 research and innovation programme (grant agreement No. 864203), BFU2017-86471-P (MINECO/FEDER, UE), “Unidad de Excelencia María de Maeztu”, funded by the AEI (CEX2018-000792-M), Howard Hughes International Early Career, Obra Social “La Caixa” and Secretaria d’Universitats i Recerca and CERCA Programme del Departament d’Economia i Coneixement de la Generalitat de Catalunya (GRC 2017 SGR 880).

## References

Auton A et al. 2015. A global reference for human genetic variation. Nature. 526:68–74. doi: 10.1038/nature15393.

Besenbacher S, Hvilsom C, Marques-Bonet T, Mailund T, Schierup MH. 2019. Direct estimation of mutations in great apes reconciles phylogenetic dating. Nat. Ecol. Evol. 3: 286–292. doi: 10.1038/s41559-018-0778-x.

Boesch C, Boesch H. 1989. Hunting behavior of wild chimpanzees in the Taï National Park. Am. J. Phys. Anthropol. 78: 547–573. doi: https://doi.org/10.1002/ajpa.1330780410.

Brawand D et al. 2011. The evolution of gene expression levels in mammalian organs. Nature. 478. doi: 10.1038/nature10532.

Fontsere C et al. 2020. Maximizing the acquisition of unique reads in non-invasive capture sequencing experiments. Mol. Ecol. Resour. 1755–0998.13300. doi: 10.1111/1755-0998.13300.

Hernandez-Rodriguez J et al. 2017. The impact of endogenous content, replicates and pooling on genome capture from faecal samples. Mol. Ecol. Resour. doi: 10.1111/1755-0998.12728.

Kaessmann H, Pääbo S. 2002. The genetical history of humans and the great apes. J. Intern. Med. 251: 1–18. doi: 10.1046/j.1365-2796.2002.00907.x.

Kim D et al. 2013. TopHat2: accurate alignment of transcriptomes in the presence of insertions, deletions and gene fusions. Genome Biol. 14:R36. doi: 10.1186/gb-2013-14-4-r36.

Kuhlwilm M et al. 2016. Evolution and demography of the great apes. Curr. Opin. Genet. Dev. doi: 10.1016/j.gde.2016.09.005.

Lawrence M, Gentleman R, Carey V. 2009. rtracklayer: an R package for interfacing with genome browsers. Bioinformatics. 25: 1841–1842. doi: 10.1093/bioinformatics/btp328.

Li H et al. 2009. The Sequence Alignment/Map format and SAMtools. Bioinformatics. 25: 2078–2079. doi: 10.1093/bioinformatics/btp352.

Li H, Durbin R. 2009. Fast and accurate short read alignment with Burrows-Wheeler transform. Bioinformatics. 25: 1754–1760. doi: 10.1093/bioinformatics/btp324.

Locke DP et al. 2011. Comparative and demographic analysis of orang-utan genomes. Nature. 469: 529–33. doi: 10.1038/nature09687.

Mallick S et al. 2016. The Simons genome diversity project: 300 genomes from 142 diverse populations. Nature. 538. doi: 10.1038/nature18964.

De Manuel M et al. 2016. Chimpanzee genomic diversity reveals ancient admixture with bonobos. Science (80-.). 354. doi: 10.1126/science.aag2602.

McKenna A et al. 2010. The Genome Analysis Toolkit: A MapReduce framework for analyzing next-generation DNA sequencing data. Genome Res. 20: 1297–1303. doi: 10.1101/gr.107524.110.

Meyer M et al. 2016. Nuclear DNA sequences from the Middle Pleistocene Sima de los Huesos hominins. Nature. 1–15. doi: 10.1038/nature17405.

Nater A et al. 2017. Morphometric, Behavioral, and Genomic Evidence for a New Orangutan Species. Curr. Biol. 27: 3487–3498.e10. doi: 10.1016/j.cub.2017.09.047.

Peyrégne S, Prüfer K. 2020. Present-Day DNA Contamination in Ancient DNA Datasets. BioEssays. 42:2000081. doi: https://doi.org/10.1002/bies.202000081.

Prado-Martinez J et al. 2013. Great ape genetic diversity and population history. Nature. 499: 471–5. doi: 10.1038/nature12228.

Prüfer K et al. 2012. The bonobo genome compared with the chimpanzee and human genomes. Nature. 486:527. doi: 10.1038/nature11128.

Prüfer K et al. 2014. The complete genome sequence of a Neanderthal from the Altai Mountains. Nature. 505: 43–9. doi: 10.1038/nature12886.

R Core Team. 2015. R: A Language and Environment for Statistical Computing. http://www.r-project.org/.

Schubert M, Lindgreen S, Orlando L. 2016. AdapterRemoval v2: rapid adapter trimming, identification, and read merging. BMC Res. Notes. 9:88. doi: 10.1186/s13104-016-1900-2.

van der Valk T, Díez-del-Molino D, Marques-Bonet T, Guschanski K, Dalén L. 2019. Historical Genomes Reveal the Genomic Consequences of Recent Population Decline in Eastern Gorillas. Curr. Biol. 29: 165–170.e6. doi: 10.1016/j.cub.2018.11.055.

White LC et al. 2019. A roadmap for high-throughput sequencing studies of wild animal populations using noninvasive samples and hybridization capture. Mol. Ecol. Resour. 19: 609–622. doi: https://doi.org/10.1111/1755-0998.12993.

Xue Y et al. 2015. Mountain gorilla genomes reveal the impact of long-term population decline and inbreeding. Science (80-.). 348: 242–245. doi: 10.1126/science.aaa3952.

